# Contrast-enhanced ultrasound measurement of pancreatic blood flow dynamics predicts type1 diabetes therapeutic reversal in preclinical models

**DOI:** 10.1101/2020.03.03.975847

**Authors:** Vinh Pham, David G. Ramirez, Richard K.P. Benninger

## Abstract

In type 1 diabetes (T1D) immune-cell infiltration into the islets of Langerhans (insulitis) and β-cell decline occurs many years before diabetes presents. Non-invasively detecting insulitis and β-cell decline would allow diagnosis of eventual diabetes and provide a means to monitor the efficacy of therapeutic intervention. However, there is a lack of validated clinical approaches for non-invasively imaging disease progression leading to T1D. Islets have a dense microvasculature that reorganizes during diabetes. We previously demonstrated contrast-enhanced ultrasound measurements of pancreatic blood-flow dynamics could predict disease progression in T1D pre-clinical models. Here we test whether these measurements can predict successful therapeutic prevention of T1D. We performed destruction-reperfusion measurements using a small-animal ultrasound machine and size-isolated microbubbles, in NOD-scid mice receiving an adoptive transfer of diabetogenic splenocytes (AT mice). Mice received vehicle control or either of the following treatments: 1) antiCD4 to deplete CD4^+^ T cells; 2) antiCD3 to block T cell activation, 3) Verapamil to reduce β-cell apoptosis and 4) TUDCA to reduce ER stress. We compared measurements of pancreas blood-flow dynamics with subsequent progression to diabetes. In AT mice blood-flow dynamics were altered >2 weeks after splenocyte transfer. AntiCD4, antiCD3 and verapamil provided a significant delay in diabetes development. Treated AT mice with delayed or absent diabetes development showed significantly altered blood flow dynamics compared to untreated AT mice. Conversely, treated AT mice that developed diabetes, despite therapy, showed similar blood-flow dynamics to untreated AT mice. Thus, contrast-enhanced ultrasound measurement of pancreas blood-flow dynamics can predict the successful or unsuccessful delay or prevention of diabetes upon therapeutic treatments that target both immune activity or β-cell protection. This strategy may provide a clinically deployable predictive marker for disease progression and therapeutic reversal in asymptomatic T1D.

## Introduction

During type1 diabetes (T1D) progression immune infiltration into the islet of Langerhans and β-cell decline occur over many years prior to diabetes onset. This asymptomatic phase presents a window in which preventative therapeutic intervention can be made, while significant β-cell mass remains [1]. Indeed, recent clinical trials have demonstrated successful delay of diabetes when applied during this asymptomatic phase [2]. However, responses to therapeutic treatments are often heterogeneous, with only a subset of subjects showing a significant delay in β-cell decline (responders) [3]. Currently there are no clinically applied indicators to assess whether the trajectory of disease progression is reversed following therapeutic treatments. While there are some clinically applied indicators of asymptomatic T1D progression [4–6], it is unclear if these indicators would be useful for assessing therapeutic efficacy: for example islet-associated autoantibodies are not pathogenic. Thus, new methods to track T1D progression are required.

We previously demonstrated that contrast-enhanced ultrasound measurements of pancreas blood flow dynamics could reveal changes in islet blood flow dynamics that result from islet microvascular remodeling during T1D progression [7–9]. We demonstrated that these measurements could predict both the speed of diabetes progression and the success of interventions deigned to prevent T1D, using preclinical mouse models of T1D [10]. In demonstrating that this approach could predict subjects in which the underlying T1D progression was halted, prior to diabetes onset, we utilized antiCD4-mediated T-cell depletion as a proof-of-principle intervention [11]. It is unclear whether the success of viable therapies for T1D prevention could also be precited by this approach.

Here we determine the degree to which contrast-enhanced ultrasound measurements of pancreas blood flow dynamics can be used to predict the success of therapies currently being applied clinically for T1D prevention or reversal. This included predicting the action of both immunotherapies and therapies directed against the β-cell. We first confirmed results from our previous study in predicting both T1D progression and the action of antiCD4 proof-of-principle treatment. We then tested whether we could predict the success of antiCD3 [12,13], verapamil [14,15] and TUDCA [16], using an adoptive-transfer model of T1D [17].

## Results

### Measuring changes to pancreas blood flow dynamics

We previously determined that measuring pancreas blood flow dynamics could indicate underlying diabetes progression in models of T1D. We set out to confirm these findings using an adoptive transfer (AT) model of diabetes. This model shows well-defined T-cell mediated autoimmune diabetes that is more consistent and rapid than in NOD mice, allowing convenient assessment of therapeutic reversal [18]. Splenocytes from recently diabetic NOD mice were transferred to NOD-scid mice. Measurements of pancreas blood flow dynamics were made prior to adoptive transfer, and at 2 weeks and 4 weeks post-transfer (**Fig.1A**). To determine whether the impact of therapeutic treatment can be predicted, different treatments were applied at 2 weeks post-transfer. At each imaging time point we performed a ‘destruction-reperfusion’ measurement whereby the reentry of blood-borne microbubble contrast agents, following high mechanical-index destruction, was measured in the pancreas by subharmonic contrast enhanced ultrasound (**Fig.1B**). We fitted the resultant contrast signal time-course following destruction to determine the rate of recovery (reperfusion rate) that can be related to perfusion speed, and the amplitude of recovery (reperfusion amplitude) that can be related to the perfusion volume [19] (**Fig.1C**).

**Figure 1.**
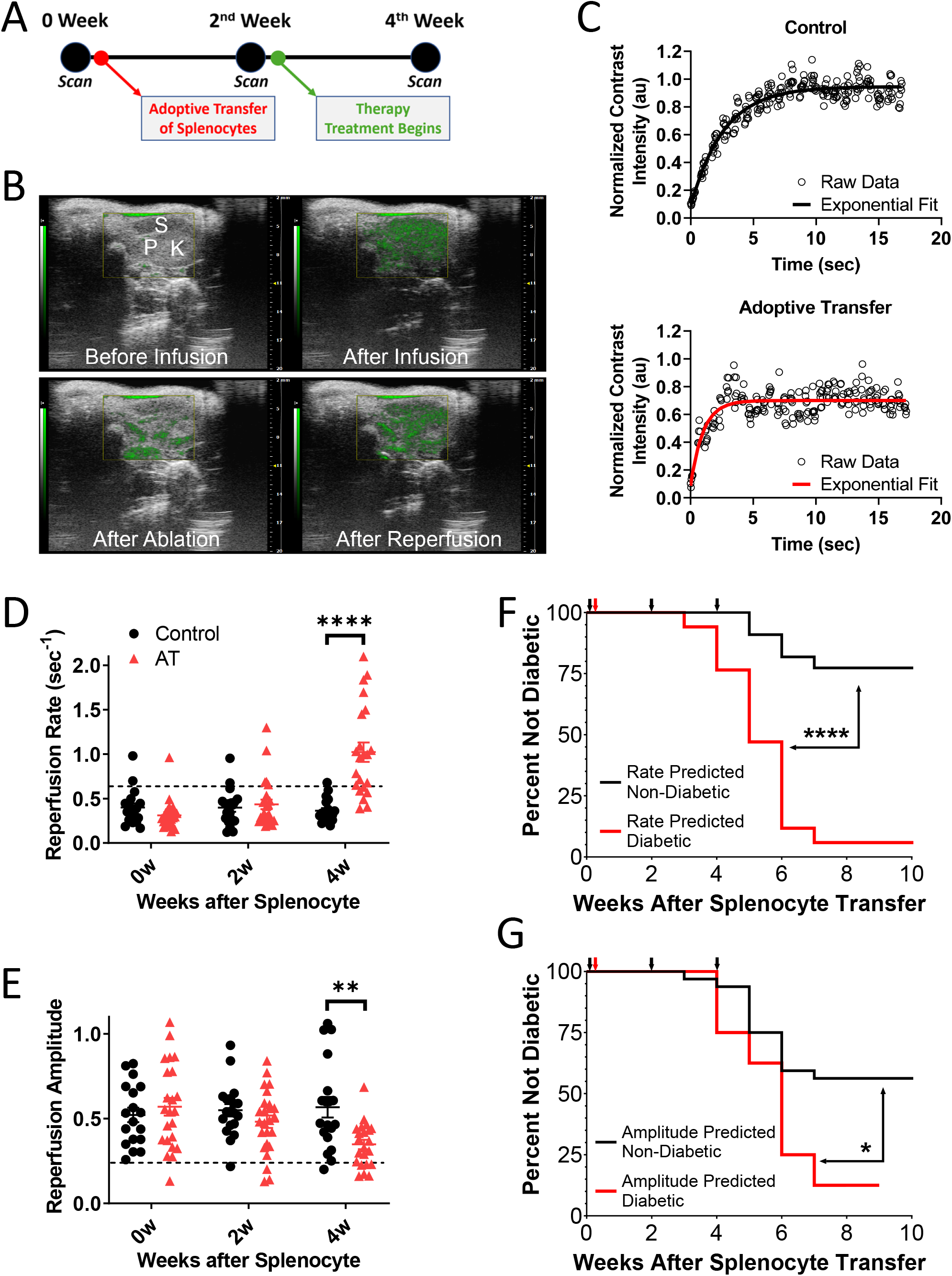
Overview of ultrasound measurements. A) Schematic of therapeutic treatments and scan times for the adoptive transfer mouse model. B) Representative images of contrast signal (green) overlaid B-mode ultrasound image of pancreas (P), kidney (K) and spleen (S), before and after microbubble infusion, and immediately following flash destruction and reperfusion. C) Representative time course of contrast signal reperfusion following flash destruction, for a control (above) and AT mouse (below) at 4 weeks following splenocyte transfer. *D*) Reperfusion rate in control animals (black) and AT animals (red) at baseline (week 0) two weeks and four weeks post splenocyte or vehicle transfer, for all animals within this study. E) as in A for reperfusion amplitude. F) Survival curves indicating diabetes development of all untreated AT mice based on measurements of the reperfusion rate at 4 weeks after splenocyte transfer. G) As in C for reperfusion amplitude at 4 weeks after splenocyte transfer. Dashed line in D,E represent disease prediction threshold that optimally separates control and untreated AT mice at age 4 weeks. * represents p<0.05, ** represents p<0.01, **** represents p<0.0001 between groups indicated, as assed by 2-sided Students t-test (D,E) or Mantel-Cox Logrank test (F,G).

We first examined the reperfusion kinetics over all control mice (lacking splenocyte transfer) and all untreated AT mice (AT mice that do not receive therapeutic drug) examined throughout this study. Consistent with prior studies, the reperfusion rate progressively increased following splenocyte transfer, and the reperfusion amplitude slightly decreased (**Fig.1D,E**). Furthermore, those animals that showed a greater increase in the reperfusion rate developed diabetes sooner than those animals that showed lower or absent increase in the reperfusion rate (**Fig.1E**). Those animals that showed a greater decrease in the reperfusion amplitude also developed diabetes sooner than those that showed lower or absent decrease in the reperfusion amplitude (**Fig.1F**), albeit with reduced significance and separation. Therefore, the reperfusion rate and to a lesser degree the reperfusion amplitude is predictive of diabetes progression in the AT model of T1D; as we previously demonstrated for the NOD model [10].

### Detecting T1D remission achieved by CD4+ T cell depletion

We previously determined that measuring the reperfusion rate could predict successful delay or prevention of diabetes in the AT model of diabetes induced by antiCD4-mediated CD4^+^ T cell depletion [10]. To confirm these findings, we repeated these experiments whereby AT mice were treated at 2 weeks of age with a single dose of antiCD4. This treatment prevented/delayed diabetes in ~50% of AT animals (**Fig.2A**). While untreated AT mice showed a significant increase in reperfusion rate, this increase was not significant following antiCD4, with a higher spread in measurements (**Fig.2B**). While consistent trends were also observed with the amplitude measurement, no significant differences were observed (**Fig.2C**). Those AT mice that showed a delay or prevention of diabetes (responders) showed a significantly lower reperfusion rate following treatment (**Fig.2D**). Conversely, those antiCD4-treated AT mice that developed diabetes at a similar time to untreated AT mice (nonresponders) showed increased reperfusion rate, similar to that in untreated animals. While trends between responders and non-responders were present with the amplitude measurement, no significant differences were observed (**Fig.2E**). To test whether the reperfusion rate measurement could predict diabetes prevention, we separated mice into two groups based on whether they showed a low reperfusion rate (similar to in control mice, predicted to be disease negative) or a high reperfusion rate (above that measured in control mice, predicted to be disease positive) – see Methods. Those treated AT animals that showed a lower reperfusion rate (predicted to be disease negative) developed diabetes with a significantly delayed and lower incidence compared to both untreated animals and to those AT animals with a higher reperfusion rate (predicted to be disease positive) (**Fig.2F**). Those animals with a higher reperfusion rate developed diabetes with a similar incidence and speed to untreated AT mice. Thus, the reperfusion rate is also predictive of successful or unsuccessful antiCD4-mediated prevention of diabetes in the AT model of T1D.

**Figure 2.**
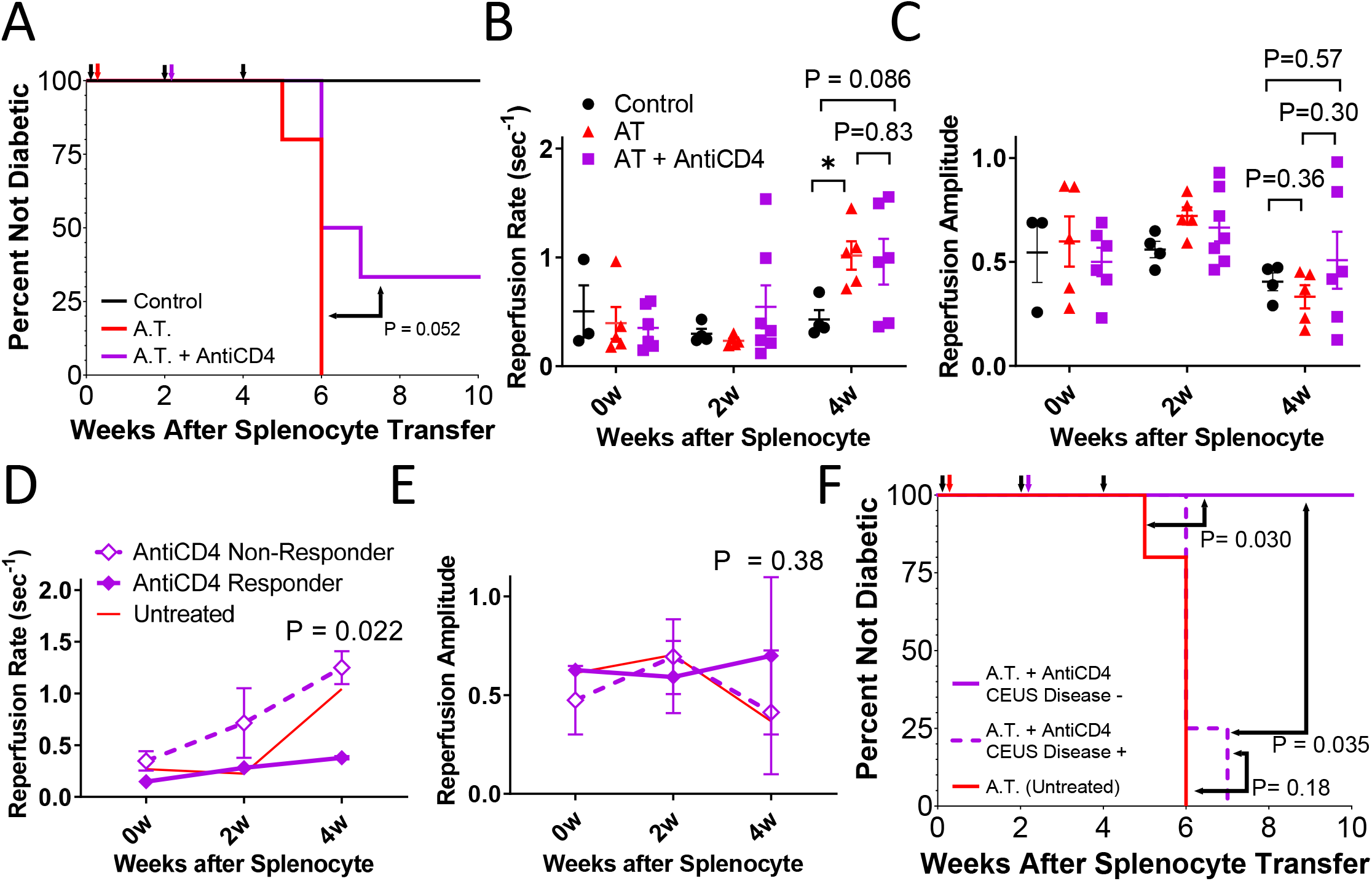
Validating the prediction of antiCD4 treatment. A) Survival curves indicating diabetes development for control (black), AT (red), and AT+anti-CD4 (purple). B) Reperfusion rate in control animals (black), AT animals (red) or AT+anti-CD4 animals (purple) at baseline (week 0) two weeks and four weeks post splenocyte or vehicle transfer. C) as in B for reperfusion amplitude. D) Average reperfusion rate in anti-CD4 responders (solid diamonds) and non-responders (open diamonds) before (week 0), two and four weeks post splenocyte transfer, together with measurement in untreated AT mice (red). E) As in D for reperfusion amplitude. F) Survival curves indicating diabetes development of untreated AT mice (solid red), together with AT+anti-CD4 mice that are predicted to not respond to anti-CD4 treatment and develop diabetes (Disease +, dashed purple) or that are predicted respond to antiCD4 treatment and not develop diabetes (Disease -, solid purple), based on measurements of the reperfusion rate at 4 weeks after splenocyte transfer. * represents p<0.05, otherwise is stated between two groups indicated, as assed by 2-sided Students t-test (B-E) or Mantel-Cox Logrank test (F).

### Detecting T1D remission achieved by antiCD3 therapy

While we could predict disease progression upon CD4+ T cell depletion, antiCD4 is not a viable therapy for T1D. We next tested whether we could predict prevention or delay of T1D upon an immunotherapy, antiCD3, that has been demonstrated to successfully delay β-cell decline at onset and prevent T1D prior to onset in clinical studies [2,12,13]. AT mice were treated at 2 weeks of age with 5 daily doses antiCD3 (clone 145-2C11). This treatment prevented/delayed diabetes in ~60% of AT animals (**Fig.3A**). On average, both untreated AT mice and antiCD3 treated mice showed a significant increase in reperfusion rate (**Fig.3B**). Similarly, both untreated AT mice and antiCD3 treated mice showed a significant decrease in reperfusion amplitude (**Fig.3C**). Those antiCD3-treated AT mice that showed a delay or prevention of diabetes (responders) showed a significantly lower reperfusion rate following treatment (**Fig.3D**). Conversely, those antiCD3-treated AT mice that developed diabetes at a similar time to untreated AT mice (non-responders) showed increased reperfusion rate, similar to that in untreated animals. As under antiCD4 treatment, no significant differences were observed in amplitude measurements between responders and non-responders (**Fig.3E**). We again separated mice into two groups based on whether they showed a low reperfusion rate (predicted to be disease negative) or a high reperfusion rate (predicted to be disease positive), Consistent with the above observations, those treated AT animals that showed a lower reperfusion rate developed diabetes with a significantly delayed and lower incidence compared to both untreated animals and AT animals with a higher reperfusion rate (**Fig.3F**). Those animals with a higher reperfusion rate developed diabetes with a similar incidence and speed to untreated AT mice. Thus, the reperfusion rate is predictive of successful or unsuccessful antiCD3-mediated therapeutic delay or prevention of diabetes in the AT model of T1D.

**Figure 3.**
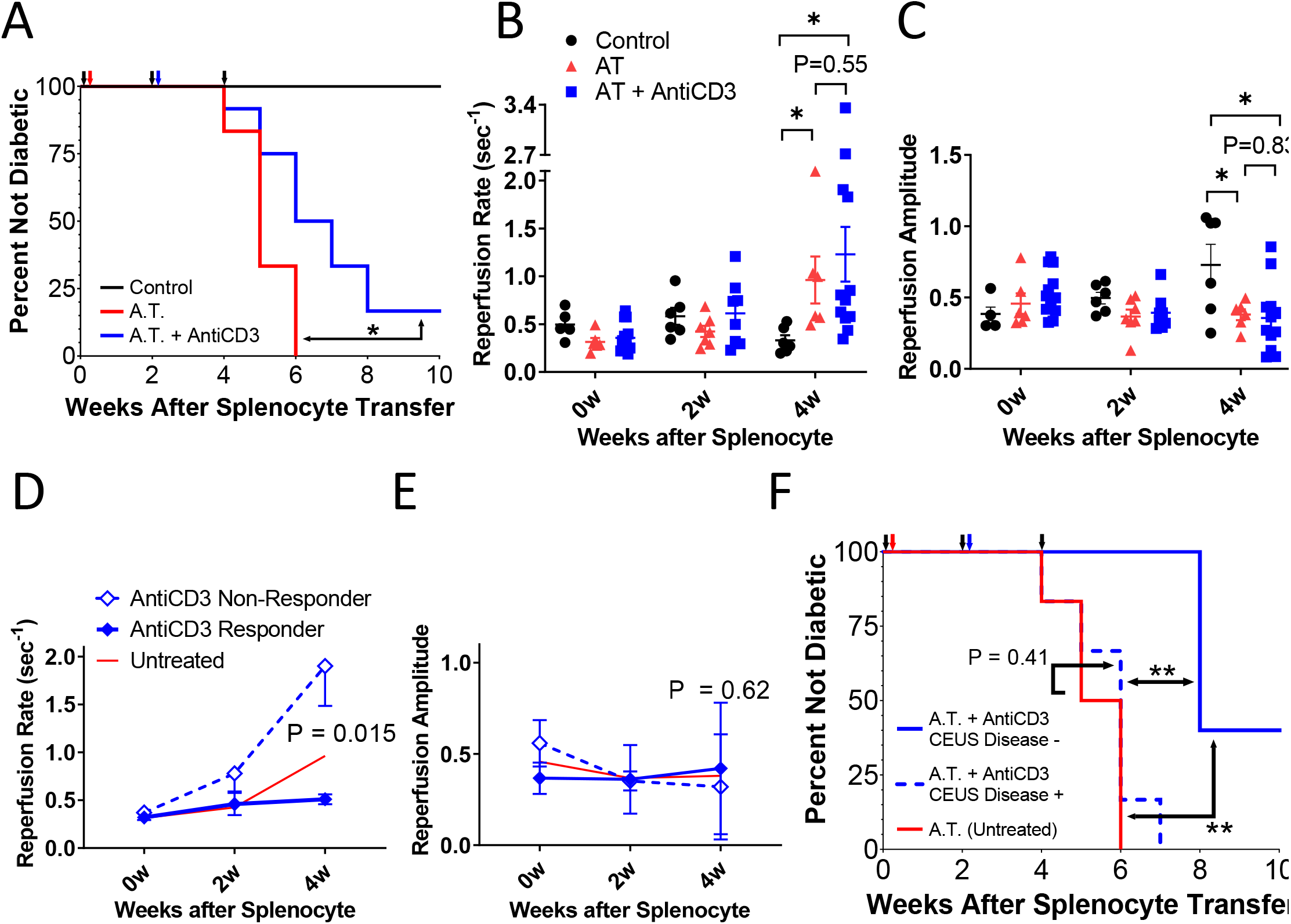
predicting antiCD3-mediated diabetes prevention. A) Survival curves indicating diabetes development for control (black), AT (red), and AT+anti-CD3 (blue). B) Reperfusion rate in control animals (black), AT animals (red) or AT+anti-CD3 animals (blue) at baseline (week 0) two weeks and four weeks post splenocyte or vehicle transfer. C) as in B for reperfusion amplitude. D) Average reperfusion rate in anti-CD3 responders (solid diamonds) and non-responders (open diamonds) before (week 0), two and four weeks post splenocyte transfer, together with measurement in untreated AT mice (red). E) As in D for reperfusion amplitude. F) Survival curves indicating diabetes development of untreated AT mice (solid red), together with AT+anti-CD3 mice that are predicted to not respond to anti-CD3 treatment and develop diabetes (Disease +, dashed blue) or that are predicted respond to anti-CD3 treatment and not develop diabetes (Disease -, solid blue), based on measurements of the reperfusion rate at 4 weeks after splenocyte transfer. * represents p<0.05, ** represents p<0.01, otherwise is stated, between two groups indicated, as assed by 2-sided Students t-test (B-E) or Mantel-Cox Logrank test (F).

### Detecting T1D remission achieved by β-cell therapies

An emerging strategy for T1D prevention is to target mechanisms underlying β-cell death/dysfunction [15,16]. We next tested whether we could predict delay or prevention of T1D upon a β-cell directed therapy, verapamil, that has been demonstrated to successfully delay β-cell decline at disease onset in clinical studies. AT mice were continually treated at 2 weeks of age onwards with verapamil in drinking water [15]. This treatment prevented/delayed diabetes in ~40% of AT animals (**Fig.4A**). Untreated AT mice showed a significant increase in reperfusion rate, but this was not significant, on average, in verapamil-treated AT mice (**Fig.4B**). Similarly, untreated AT mice showed a significant decrease in reperfusion amplitude, but this was not significant, on average, in verapamil-treated AT mice (**Fig.4C**). Those verapamil-treated AT mice that showed a delay or prevention of diabetes (responders) showed a significantly lower reperfusion rate following treatment (**Fig.4D**). Conversely, those verapamil-treated AT mice that developed diabetes at a similar time to untreated AT mice (nonresponders) showed increased reperfusion rate, similar to that in untreated animals. As under antiCD4 or antiCD3 treatment, no significant differences were observed in amplitude measurements between responders and non-responders (**Fig.4E**). Upon separating mice into two groups, with low reperfusion rate (disease negative) or a high reperfusion rate (disease positive), those verapamil-treated AT animals that showed a lower reperfusion rate developed diabetes with a significantly delayed and lower incidence than those with a higher reperfusion rate (**Fig.4F**). Those treated animals with a higher reperfusion rate developed diabetes with a similar incidence and speed to untreated AT mice. Thus, the reperfusion rate is predictive of successful or unsuccessful verapamil-mediated therapeutic prevention of diabetes in the AT model of T1D.

**Figure 4.**
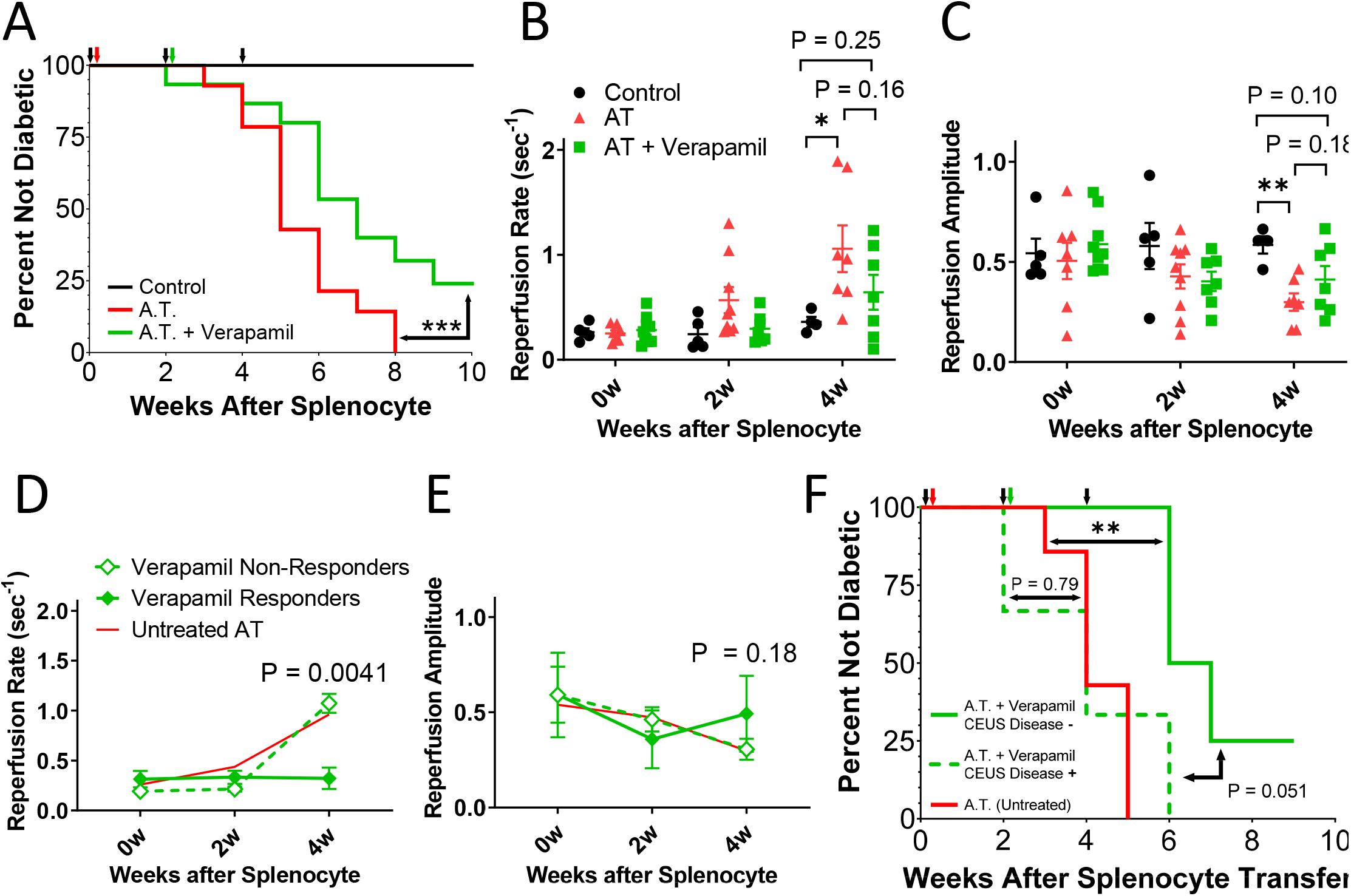
predicting verapamil-mediated diabetes prevention. A) Survival curves indicating diabetes development for control (black), AT (red), and AT+verapamil (green). B) Reperfusion rate in control animals (black), AT animals (red) or AT+ verapamil animals (green) at baseline (week 0) two weeks and four weeks post splenocyte or vehicle transfer. (C) as in B for reperfusion amplitude. (D) Average reperfusion rate in verapamil responders (solid diamonds) and non-responders (open diamonds) before (week 0), two and four weeks post splenocyte transfer, together with measurement in untreated AT mice (red). E) As in D for reperfusion amplitude. F) Survival curves indicating diabetes development of untreated AT mice (solid red), together with AT+verapamil mice that are predicted to not respond to verapamil treatment and develop diabetes (Disease +, dashed green) or that are predicted respond to verapamil treatment and not develop diabetes (Disease -, solid green), based on measurements of the reperfusion rate at 4 weeks after splenocyte transfer. * represents p<0.05, ** represents p<0.01, otherwise is stated, between two groups indicated, as assed by 2-sided Students t-test (B-E) or Mantel-Cox Logrank test (F).

Finally, we examined another β-cell directed therapy, Tauroursodeoxycholic Acid (TUDCA), that is being applied in clinical studies (ClinicalTrials.gov Identifier: NCT02218619). AT mice were continually treated at 2 weeks of age onwards with TUDCA via IP injection [16]. In contrast to verapamil treatment, we did not observe a significant impact on diabetes development by TUDCA (**SI Fig.S1A,B**). Both untreated AT mice and TUDCA-treated AT mice showed a significant increase in reperfusion rate, with no difference between these groups (**SI Fig.S1C**). Similarly, no difference in reperfusion amplitude was observed between untreated AT mice and TUDCA-treated AT mice (**SI Fig.S1D**). Thus the reperfusion rate was predictive of the lack of efficacy, in our hands, by TUDCA.

## Discussion

In this study we sought to test whether changes in contrast-enhanced ultrasound measurement of pancreas blood flow dynamics could be used to predict the therapeutic prevention of type1 diabetes (T1D). We demonstrated that the delay and prevention of diabetes induced by antiCD4-mediated depletion of CD4^+^ T cells and antiCD3-mediated blocking of T cell activation could be predicted in an adoptive transfer model of T1D. Furthermore, we demonstrated that the delay/prevention of diabetes induced by verapamil-mediated protection of β-cell decline could also be predicted in an adoptive transfer model of T1D. Under conditions where no diabetes was delayed/prevented, during TUDCA treatment, no difference was observed, strengthening how the measurement of pancreas blood flow dynamics strongly correlated with diabetes progression.

Our data suggests that contrast-enhanced ultrasound measurement of pancreas blood flow dynamics could be a viable approach to predict successful prevention of diabetes when treated during the asymptomatic phase of T1D. This is critically important, given heterogeneous or incomplete responses to therapeutic reversal [3] or prevention [2] of T1D in human trials and the lack of any approach presently that can predict the therapeutic reversal of T1D [20]. While autoantibodies can be used to predict onset of diabetes [4], it is unclear whether they can be used to predict therapeutic prevention. We further show this approach is applicable to therapies that have been demonstrated to modulate autoimmunity or β-cell viability, indicating it is broadly applicable.

It is unclear what the specific mechanism is that links T1D progression and pancreas blood flow dynamics. We and others have previously demonstrated in models of T1D that changes to the pancreas blood flow reflect changes to islet-specific blood flow dynamics [10,21,22]. Islet blood flow dynamics also correlate with remodeling of the islet microvasculature that occurs in both mouse models of T1D and human T1D [7–9,23,24]. Altered microvascular organization may result from inflammation and thus lead to altered islet blood flow [7,25,26]. However, altered pancreas blood flow, for example as a result of islet decline or upstream arterial blood flow could also lead to islet microvascular reorganization [22,27]. While we have demonstrated predictive power for pancreas blood flow dynamics, determining the mechanism underlying altered blood flow dynamics in T1D will be important to fully interpret this diagnostic measurement. For example, the factor associated with T1D that determines blood flow dynamics may not fully explain T1D progression or reversal. This may include important differences between mouse models and human T1D.

The approach we use here is applicable to human subjects: contrast-enhanced ultrasound has been applied, albeit off label, to pancreas imaging for other indications [28–30]. The islet microvasculature shows remodeling in human T1D, although studies prior to diabetes onset are lacking. However, the mouse and human islet do show important differences in terms of islet blood flow and microvasculature organization [31–33]. Some therapies such as antiCD3 may also show differing mechanism of action between human and mouse [34]. Therefore, testing this approach in human T1D or with the aid of humanized models for T1D are still needed.

In summary, we demonstrate that ultrasound measurement of pancreas blood flow dynamics can predict successful therapeutic prevention of diabetes, induced by therapies relevant to treating human T1D. This potentially provides a method to translate to human studies, in which no approach currently exists to assess the therapeutic prevention of T1D.

## Methods

### Animals

All animal procedures were performed in accordance with protocols approved by the Institutional Animal Care and Use Committee of the University of Colorado Anschutz Medical campus. Female NOD mice were purchased from Jackson Laboratories (Bar Harbor, ME) at age 4 weeks. Female NOD-SCID animals were purchased from Jackson Laboratories at age 10-14 weeks. Throughout the study, animals were monitored weekly for blood glucose concentration utilizing a blood glucometer (Bayer).

### Isolation and Adoptive Transfer of Diabetogenic Splenocytes

Splenocytes were isolated from diabetic female NOD mice (hyperglycemic <1 week), manually dissociated and counted in cold HBSS (without MgCl_2_ and CaCl_2_). Leukocytes were counted to determine an estimate of cellular density. NOD-SCID mice received a single intraperitoneal (I.P.) dose of 20 x 10^6^ leukocytes resuspended in HBSS. Control animals were injected with an equivalent volume of HBSS without leukocytes.

### Contrast-Enhanced Ultrasound (CEUS) imaging

General anesthesia was established with isoflurane inhalation for a total of 20-25 minutes for all animal imaged. Prior to imaging, a custom made 27G ½” winged infusion set (Terumo BCT, Lakewood, CO) was attached to a section of polyethylene tubing (0.61 OD x 0.28 ID; PE-10, Warner Instruments) and was inserted in the lateral tail vein and secured with VetBond (3M). Abdominal fur was removed using depilatory cream, and ultrasound coupling gel placed between the skin and transducer. Foot pad electrodes on the ultrasound machine platform monitored the animal’s electrocardiogram, respiration rate, and body temperature. All animals were constantly monitored throughout the imaging session to maintain body temperature and respiration rate.

A VEVO 2100 small animal high-frequency ultrasound machine (Visual Sonics, Fujifilm, Toronto, Canada) was used for all experiments. For CEUS imaging a MS250 linear array transducer was used at a frequency of 18 MHz. B-mode imaging (transmit power 100%) was performed prior to NB or MB infusion to identify anatomy of the pancreas body, based on striated texture and location in relation to the spleen, kidney, and stomach [10]. Following identification of the pancreas and selection of a region of interest, sub-harmonic contrast mode was initiated. Acquisition settings were set at: transmit power 10%, (MI=0.12), frequency 18 MHz, standard beamwidth, contrast gain of 30 dB, 2D gain of 18 dB, with an acquisition rate of 26 frames per second.

Size-isolated microbubble contrast agent (‘SIMB3-4’, Advanced Microbubble Laboratories) was injected as a single bolus of ~10 million bubbles in phosphate buffered saline (pH 7.4) into the lateral tail vein via the catheter. SIMBs were allowed to circulate throughout the animal for ~20 seconds to reach a relative steady state of systemic distribution. SIMB destruction was initiated by delivery of a high mechanical index pulse (VEVO2100 burst mode, MI=0.2), to destroy a portion of SIMBs within the imaging plane [35,36]. Data were acquired for at least 10 seconds following SIMB destruction to adequately measure reperfusion into the tissue.

### Data Analysis

Gating to remove movements as a result of animal breathing was carried out manually or in MATLAB (MathWorks, Natick, MA). For analysis of reperfusion kinetics, the background non-linear intensity taken before SIMB infusion was subtracted from the entire trace. Each reperfusion time-course was normalized to a 0.5s average of the steady state NL contrast intensity immediately prior to flash destruction. The resultant normalized reperfusion curves were fit in MATLAB with an exponential rise equation, *F*(*t*) = *C* + *A*(1-e^-*kt*^), where *C* is the offset from zero, *A* is the amplitude of the curve and *k* is the rate of reperfusion. CEUS measurements were excluded from analysis if a poor infusion was recorded and/or poor microbubble flash-destruction occurred (and thus poorly defined recovery).

### Therapeutic drug application

For antiCD4 treatment, NOD-Scid animals two weeks following splenocyte delivery were injected with a single dose of 20 mg anti-mouse CD4, clone GK1.5 (BP0003-1; BioXCell, W. Lebanon, NH). For antiCD3 treatment, NOD-Scid animals two weeks following splenocyte delivery were injected with 5 daily doses of 5μg anti-mouse CD3, clone 145-2C11 (BP0001-1; BioXCell). For verapamil treatment, NOD-Scid animals two weeks following splenocyte delivery received verapamil (Thermo Fisher Scientific, Waltham MA) in the drinking water (1 mg/ml), available continuously ad lib. For TUDCA treatment, NOD-Scid animals two weeks following splenocyte delivery were injected daily with 300mg/kg body weight TUDCA (Millipore Sigma, Burlington MA) dissolved in sterile PBS for 14 days.

### Statistical Analysis

All data are presented as means ± SEM. Statistical comparisons were made using paired or unpaired student’s t-tests or ANOVAs where appropriate and as indicated. Outliers, as defined by Grubbs’ test, were excluded for the purpose of statistical analysis but retained in figure panels. Statistical significance was taken as p<0.05.

When testing for disease prediction, Receive Operating characteristics (ROC) curves were generated where sensitivity was defined as % of AT mice within some disease threshold and 1-specificty was defined as % of vehicle control mice within some disease threshold. This was performed for all Control and untreated AT mice (**Fig.1d-g**). Maximum likelihood analysis was employed to define an optimum disease prediction threshold. For each group of treated AT mice, this threshold was then employed to separate treated AT mice at 4 weeks post-transfer into disease positive and disease negative groups, which were subsequently compared.

Sample sizes for experimental groups were based on our prior study [10], to provide sufficient statistical power given the measured effect size (reperfusion parameters). When comparing experimental groups, CEUS recordings were not made in a defined order.

## Acknowledgements

Richard KP Benninger is the guarantor of this work and, as such, had full access to all the data in the study and takes responsibility for the integrity of the data and the accuracy of the data analysis. All authors acknowledge that no conflict of interest exists. This work was supported by Juvenile Diabetes Research Foundation Grants 1-INO-2017-435-A-N, 1-INO-2019-783-A-B, 5-CDA-2014-198-A-N; and NIH grants R01 DK102950, R01 DK106412 (to RKPB). DR has received funding from NIH training grant T32 HL072738-14 and F31 DK121488; and NSF Grant HRD-1301885 (sub-award, G-8960-1). The funders had no role in the study design, data collection and analysis, decisions to publish, or preparation of the manuscript.

## Author Contributions

VP designed and performed experiments, analyzed data, and edited the manuscript; DR performed experiments and analyzed data; RKPB conceived of the idea, designed experiments and wrote the manuscript.

## Supplementary Information

**Figure S1.**
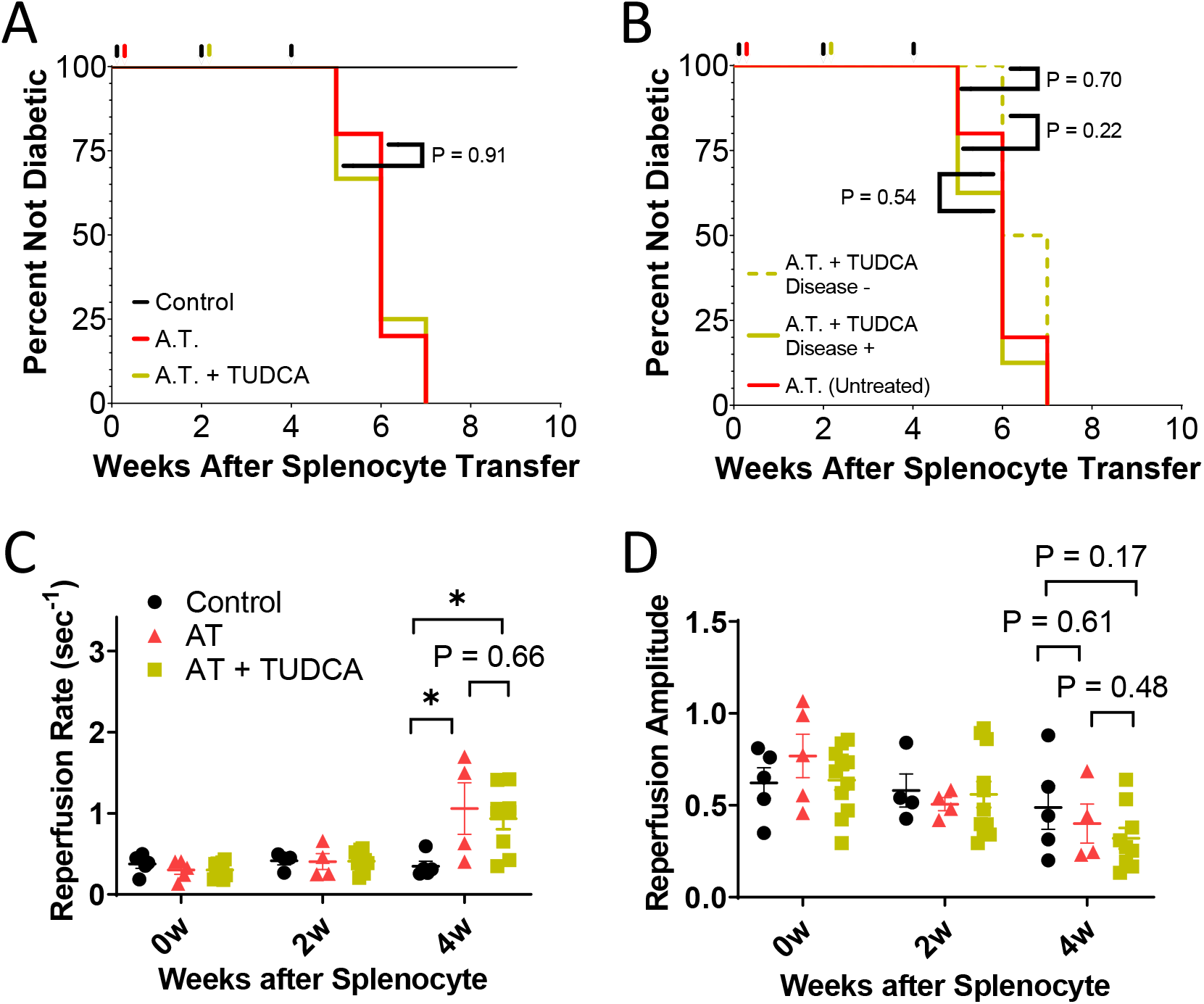
predicting TUDCA-mediated diabetes prevention. A) Survival curves indicating diabetes development for control (black), AT (red), and AT+TUDCA (yellow). B) Survival curves indicating diabetes development of untreated AT mice (solid red), together with AT+TUDCA mice that are predicted to not respond to TUDCA treatment and develop diabetes (Disease +, dashed yellow) or that are predicted respond to TUDCA treatment and not develop diabetes (Disease -, solid yellow), based on measurements of the reperfusion rate at 4 weeks after splenocyte transfer. C) Reperfusion rate in control animals (black), AT animals (red) or AT+TUDCA animals (yellow) at baseline (week 0) two weeks and four weeks post splenocyte or vehicle transfer. D) as in C for reperfusion amplitude. * represents p<0.05, otherwise is stated, between two groups indicated, as assed by 2-sided Students t-test (C,D) or Mantel-Cox Logrank test (B).

**Figure S2.**
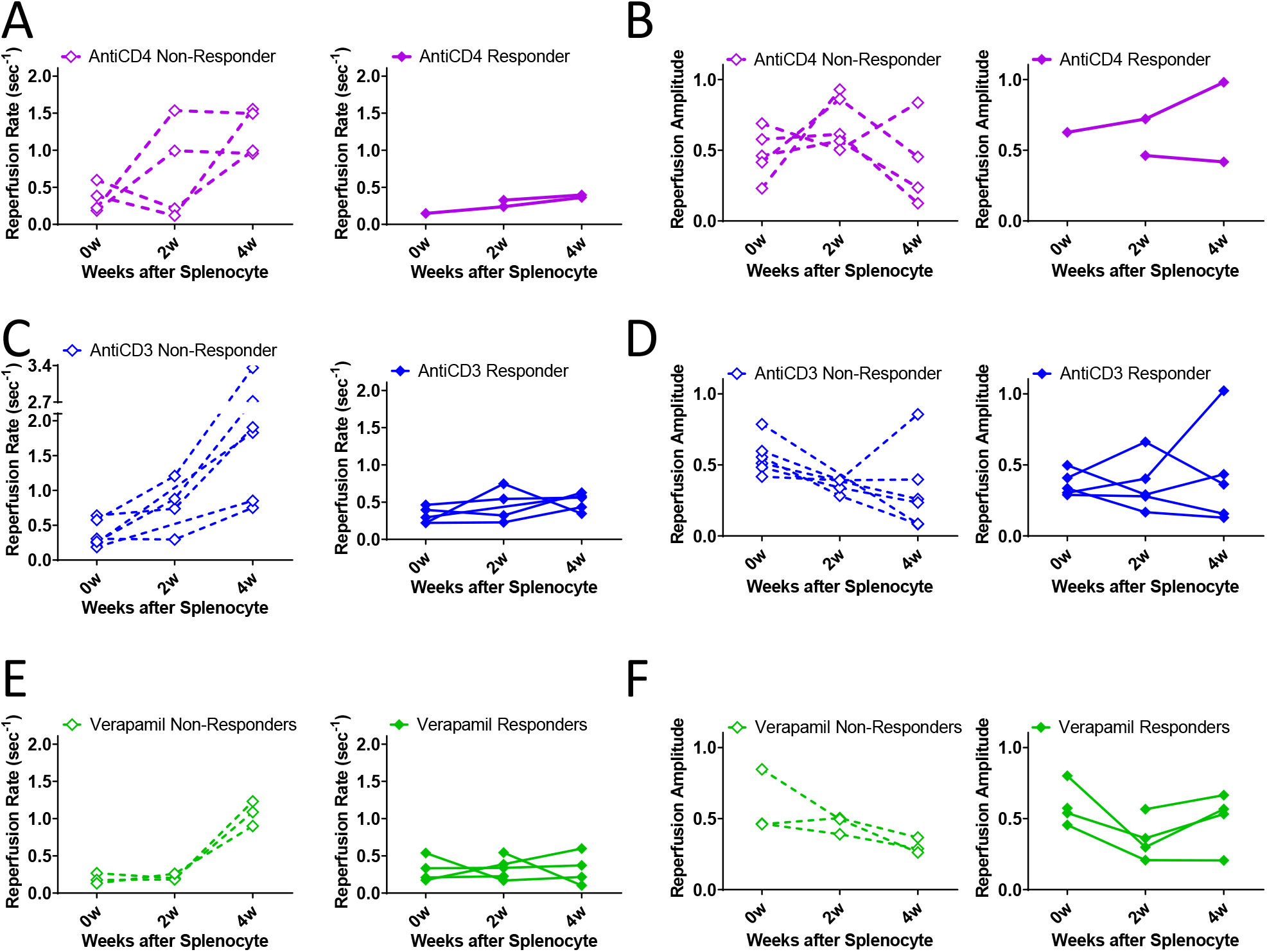
Individual reperfusion measurements for therapeutic responders and non-responders. A) Individual reperfusion rates in anti-CD4 responders (purple solid diamonds) and non-responders (purple open diamonds) before (week 0), two and four weeks post splenocyte transfer. B) as in A for reperfusion amplitude. C) Individual reperfusion rates in anti-CD3 responders (blue solid diamonds) and nonresponders (blue open diamonds) before (week 0), two and four weeks post splenocyte transfer. D) as in C for reperfusion amplitude. E) Individual reperfusion rates in verapamil responders (green solid diamonds) and non-responders (green open diamonds) before (week 0), two and four weeks post splenocyte transfer. F) as in E for reperfusion amplitude.

